# Shiftless Restricts Viral Gene Expression and Influences RNA Granule Formation during KSHV lytic replication

**DOI:** 10.1101/2022.02.18.480778

**Authors:** William Rodriguez, Timothy Mehrmann, Mandy Muller

## Abstract

Herpesviral infection reflects thousands of years of co-evolution and the constant struggle between virus and host for control of cellular gene expression. During Kaposi’s sarcoma-associated herpesvirus (KSHV) lytic replication, the virus rapidly seizes control of host gene expression machinery by triggering a massive RNA decay event *via* a virally-encoded endoribonuclease, SOX. This virus takeover strategy decimates close to 80% of cellular transcripts, reallocating host resources toward viral replication. The host cell, however, is not entirely passive in this assault on RNA stability. A small pool of host transcripts that actively evade SOX cleavage has been identified over the years. One such “escapee”, C19ORF66 (herein referred to as Shiftless - SHFL) encodes a potent anti-viral protein capable of restricting the replication of multiple DNA, RNA, and retroviruses including KSHV. Here, we show that SHFL restricts KSHV replication by targeting the expression of critical viral early genes, including the master transactivator protein, KSHV ORF50, and thus subsequently the entire lytic gene cascade. Consistent with previous reports, we found the SHFL interactome throughout KSHV infection is dominated by RNA-binding proteins that influence both translation and protein stability, including the viral protein ORF57, a crucial regulator of viral RNA fate. We next show that SHFL affects cytoplasmic RNA granule formation, triggering the disassembly of processing bodies. Taken together, our findings provide insights into the complex relationship between RNA stability, RNA granule formation, and the anti-viral response to KSHV infection.

**Significance:** In the past five years, SHFL has emerged as a novel and integral piece of the innate immune response to viral infection. SHFL has been reported to restrict the replication of multiple viruses including several flaviviruses and the retrovirus HIV-1. However, to date, the mechanism(s) by which SHFL restricts DNA virus infection remains largely unknown. We have previously shown that following its escape from KSHV-induced RNA decay, SHFL acts as a potent anti-viral factor, restricting nearly every stage of KSHV lytic replication. In this study, we set out to determine the mechanism by which SHFL restricts KSHV infection. We demonstrate that SHFL impacts all classes of KSHV genes and found that SHFL restricts the expression of several key early genes, including KSHV ORF50 and ORF57. We then mapped the interactome of SHFL during KSHV infection and found several host and viral RNA-binding proteins that all play crucial roles in regulating RNA stability and translation. Lastly, we found that SHFL expression influences RNA granule formation both outside of and within the context of KSHV infection, highlighting its broader impact on global gene expression. Collectively, our findings highlight a novel relationship between a critical piece of the anti-viral response to KSHV infection and the regulation of RNA-protein dynamics.

## Introduction

Kaposi’s sarcoma-associated herpesvirus or Human-Herpesvirus-8 (KSHV/HHV-8) is an oncogenic gamma-2-herpesvirus and the causative agent of multiple malignancies including its namesake Kaposi’s Sarcoma (KS), and two lymphoproliferative disorders: primary effusion lymphoma (PEL), and Multicentric Castleman’s disease (MCD) (**1,2**). Like all herpesviruses, KSHV infection is defined by two distinct phases: a life-long viral latency, where most of viral gene expression is suppressed, broken only by active lytic viral replication (**3**). KSHV latency establishment facilitates virus persistence within its human host for decades, reflecting KSHV’s exceptional capacity to evade detection by host immune surveillance. Sporadically, in response to a growing list of environment and intracellular triggers, KSHV switches into a lytic replicative state, swiftly remodeling and repurposing the host cell towards viral gene expression and progeny virion assembly (**3,4**). Normally, the human immune response keeps this life cycle in-check by actively suppressing KSHV infection. However, in immuno-compromised individuals such as untreated AIDS patients, KSHV infection can lead to the production of pro-tumorigenic factors (both viral and host) that drive forward KSHV-associated malignancies (**5, 6**).

Successful KSHV lytic replication relies on the ability of the virus to rapidly seize and maintain control of cellular gene expression following reactivation from latency. One KSHV stratagem for taking over these resources is to trigger a global RNA decay event termed “host-shutoff”, which is orchestrated by KSHV’s ORF37 (SOX), a virally-encoded endoribonuclease (**7–10**). SOX expression decimates most of the host transcriptome, releasing host resources, once bound to cellular mRNA, now free for coordination of viral gene expression (**10–12**).

While the breadth of mRNA targeted by SOX and other viral host-shutoff proteins is expansive, we and others have found that a select few host mRNAs are spared from degradation (**13, 14**). These transcripts, termed “escapees”, are spared from a range of viral—but not host—endonucleases (**13–18**) and appear to actively escape cleavage *via* a protective RNA element located within their 3’ UTRs that we refer to as the “SOX-resistant element” or SRE. These “dominant” escapees previously included only the host interleukin-6 (IL-6) (**16**) and the growth arrest and DNA damage-inducible 45 beta (GADD45B) (**15**). More recently, we identified yet another SRE-bearing mRNA, C19ORF66 (RyDEN, IRAV, SVA-1, Shiftless), herein referred to as SHFL, a cellular transcript that not only evades cleavage by SOX but multiple herpesviral endonucleases and even the Influenza A virus (IAV) PA-X endonuclease (**18**). Upon further investigation, we demonstrated that SHFL is a stringent anti-KSHV factor, restricting KSHV lytic reactivation from latency and all subsequent stages of lytic viral replication.

SHFL is an interferon stimulated gene (ISG) that is demonstrably a vital piece of the innate immune response to viral infection, capable of suppressing the replication of multiple DNA, RNA, and retroviruses (**18–27**). Studies of SHFL function over the past 5 years have revealed its multifaceted capacity to negatively modulate viral RNA stability, viral gene translation, and even viral protein stability through interactions with cellular co-factors that coordinate these processes such as cytoplasmic poly(A) binding protein 1 (PABPC1), La Ribonucleoprotein Domain Family Member 1 (LARP1) and the RNA helicase MOV10 (**19,20**). SHFL can also induce the degradation of viral proteins through a variety of pathways including lysosomal degradation (**22**) and ubiquitination *via* the E3 ubiquitin ligase MARCH8 (**25**). SHFL is also among the first human genes identified as capable of restricting the −1 Programmed Ribosomal Frameshift (−1PRF) of both HIV-1 and the current pandemic associated virus, SARS-CoV-2 (**21,24**). Collectively, SHFL has emerged as a critical piece of the innate host defense array of ISGs against viral infection.

While the breadth of SHFL activity continues to be unraveled, the molecular mechanism(s) behind SHFL’s function, especially during DNA virus infection, remain largely unknown. Here, we demonstrate that SHFL broadly restricts KSHV lytic gene expression, including that of the master latent-to-lytic switch protein, KSHV’s ORF50 (RTA), an impact that catastrophically dysregulates the initiation of the KSHV lytic gene cascade. Upon further investigation, we found that the SHFL interactome is dominated by RNA-binding proteins during both KSHV latency and lytic replication. Surprisingly, we found that SHFL negatively influences the assembly of a key RNP granule type, RNA Processing bodies (P-bodies), while simultaneously inducing the formation of Stress Granules (SGs) both outside of and within the context of KSHV infection. Lastly, SHFL also interacts with and restricts the expression of the KSHV RNA-binding protein ORF57, a broad regulator of viral RNA fate. Taken together, our findings highlight that SHFL is an integral piece of the virus-host arms between KSHV and its human host for control of cellular gene expression during lytic replication following its escape from SOX cleavage.

## Results

### SHFL Broadly Restricts KSHV Lytic Gene Expression

SHFL is a potent anti-viral factor capable of restricting the replication of multiple viral families including flaviviruses, alphaviruses, and retroviruses. Consistently, we have previously demonstrated that SHFL also restricts KSHV lytic replication (**18**). To determine the mechanism underlying this restriction, we first investigated the breadth of SHFL impact on KSHV lytic gene expression. First, we examined an array of viral genes spanning all kinetic classes, both at the RNA and protein levels when overexpressing SHFL in the KSHV-positive renal carcinoma cell line, iSLK.Bac16 (herein referred to as iSLK.WT). We observed that SHFL moderately to severely restricted gene expression for all lytic viral gene products tested (**Figure 1A-B, Supp Figure 1**). Given this extensive effect on lytic gene expression, we hypothesized that instead of individually targeting each of these genes, SHFL could be targeting one of the earliest and most critical regulators of the viral lytic gene cascade: KSHV’s ORF50. ORF50 (RTA) is a master viral transcriptional regulator that controls the switch between the KSHV latency and active lytic replication (**28–32**). ORF50 alone has been shown to be both necessary and sufficient for lytic reactivation and the transactivation of multiple lytic gene promoters, including its own (**29**). In line with our hypothesis, using both overexpression and knockdown approaches, we confirmed that ORF50 expression is severely restricted by SHFL (**Figure 1C-D**). Furthermore, in iSLK.WT cells, ORF50 is expressed from two distinct promoters: from the “endogenous” one on the viral genome (viral ORF50 or vORF50) and from an exogenous doxycycline-inducible promoter (exogenous or eORF50) used to artificially promote the latent-to-lytic switch (**33**). The ORF50 detected in our initial RT-qPCR screen represents the total amount of ORF50 expressed upon induction of lytic reactivation from both promoters. To determine whether SHFL effect on ORF50 was promoter-dependent, we next designed a set of primers to differentially assess vORF50 vs. eORF50 expression (**Figure 1E**). We observed that SHFL ability to repress ORF50 expression was active regardless of the ORF50 source, suggesting that SHFL-mediated effect is promoter independent. Lastly, Since SHFL has been shown to interact with viral RNAs, we next checked whether it could directly bind to ORF50 mRNA. To test this, we used RNA immunoprecipitation and found that SHFL does bind to ORF50 mRNA during KSHV lytic replication (**Figure 1F**). We thus hypothesized that SHFL-mediated repression of ORF50 could stem from a destabilization of ORF50 mRNA. However, using an Actinomycin D assay, we did not observe any significant difference in ORF50 mRNA half-life (t_1/2_ = 4h) upon SHFL expression (**Supp Fig 2**). Collectively, these data suggest that SHFL restricts lytic gene expression post-transcriptionally and in a manner independent from viral RNA stability.

**Figure 1:**
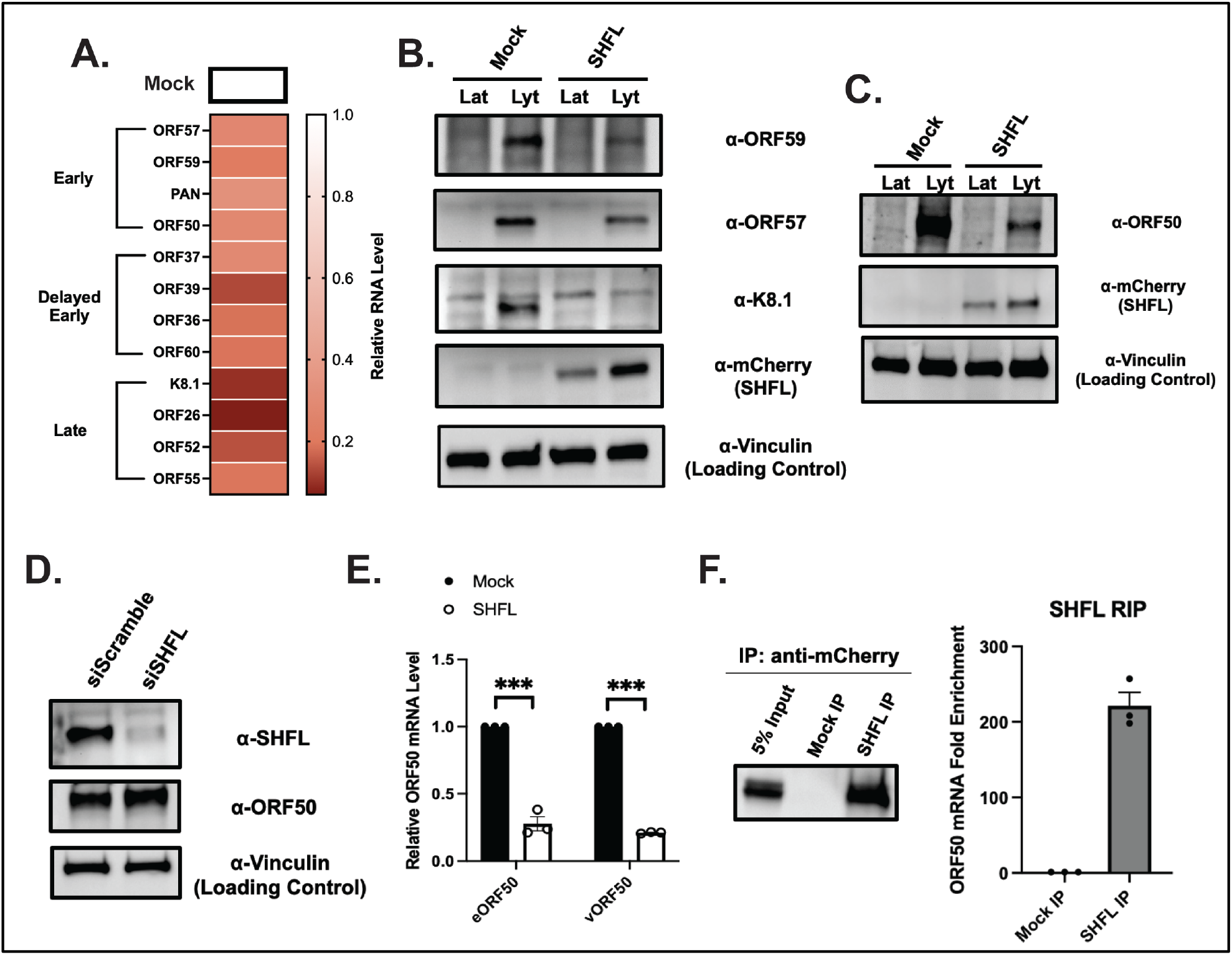
SHFL broadly restricts KSHV Lytic Gene Expression. (**A**) KSHV-positive iSLK.WT cells were transfected with a FLAG-tagged SHFL or a FLAG-empty vector and reactivated with doxycycline and sodium butyrate for 48h. Total RNA was then harvested and subjected to RT-qPCR to measure mRNA levels of the indicated viral early, delayed early, and late genes. Represented here is a heat map summarizing data from three biological replicates shown in Supplementary Figure 1. (**B, C**) iSLK.WT cells were first transfected with either a Nter-tagged mCherry-SHFL (NC-SHFL) or a mCherry only (mock) vector. Cells were then reactivated for 48h (lyt) or left untreated (lat). Cells were then harvested, lysed, resolved on SDS-PAGE, and immunoblotted with the indicated antibodies. (**D**) Unreactivated or reactivated iSLK.WT cells were treated with either siRNA targeting SHFL (or control scramble siRNAs) for 48h. siRNA-treated cells were then harvested, lysed, resolved on SDS-PAGE, and immunoblotted with the indicated antibodies. (**E**) iSLK.WT cells were transfected with either NC-SHFL or a mock vector and reactivated for 48h. (**F**) iSLK.WT cells were transfected with either NC-SHFL or a mock vector and reactivated for 48h. Cells were then harvested, lysed, and RNA immunoprecipitation (RIP) performed using mCherry antibody. Following reverse crosslinking, total RNA was then harvested and subjected to RT-qPCR using primers as indicated. Statistics were determined using students paired t-test between control and experimental groups; error bars represent standard error of the mean; n=3 independent biological replicates.

### SHFL Interacts with RNA-binding proteins during KSHV Infection

To better understand how SHFL is regulating ORF50 expression, we next set out to map its interaction network throughout KSHV infection. iSLK.WT were either left latent or were reactivated for 48h with doxycycline and sodium butyrate to trigger KSHV lytic cycle. After verifying SHFL pull-down efficiency (**Supp Fig 3**), the SHFL interactome during KSHV infection was mapped using LC-MS/MS. In total, 98 unique proteins were identified as SHFL interactors, of which 9 were exclusively detected in the latent cells and 12 were exclusively found in lytic cells (**Supp Table 1)**, (**Figure 2A**). The remaining interactors span both latency and lytic replication and include the known SHFL interactor PABPC1, which we also confirmed via co-immunoprecipitation (**Supp Fig 3**) (**19**). Gene Ontology analysis on SHFL interactors revealed several functional categories including RNA binding and Ubiquitin ligase binding, confirming SHFL previously suspected roles in both RNA and protein stability (**Figure 2B-C**) (**Supp Table 2**). Notably, several important cellular RNA binding proteins were identified that are known constituents of cytoplasmic stress granules including PABPC1, KPNA2, DDX3X, FUS, and HNRNPK (**34–37**). Intriguingly, we also detected three viral proteins as potential SHFL interactors: ORF59, the KSHV DNA processivity factor; ORF57, the master regulator of KSHV RNA fate and ORF52, a tegument protein that inhibits cytosolic viral DNA sensing via cGAS/STING (**38–40**).

**Figure 2:**
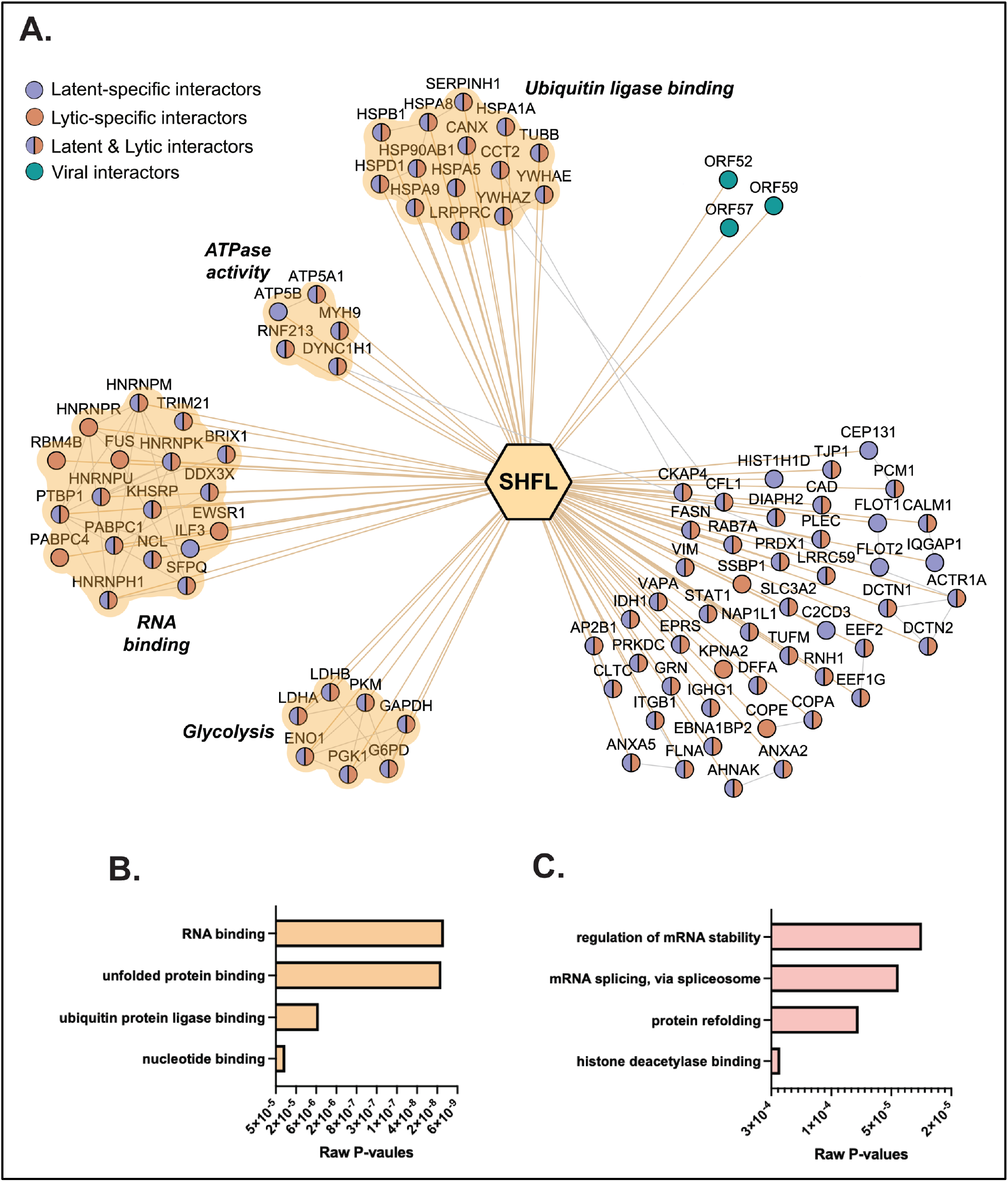
Determination of SHFL interactome during KSHV infection. (**A**) iSLK.WT cells were reactivated for 72h and subjected to mass spectrometry. Network generated by Cytoscape represents the interactome of SHFL during both KSHV latency and lytic replication. 92 high confidence interactions between SHFL (center hexagon) and human proteins (nodes) were identified by mass spectrometry along with 3 viral proteins (blue nodes). Interactions exclusively identified in latent samples are labelled in purple while those identified specifically in the lytic samples are identified in orange. Physical interactions among host proteins (thin gray lines) were manually curated from the STRING and IntAct databases. Gene Ontology (GO) enrichment analysis was performed on the human interacting proteins of SHFL using DAVID bioinformatic database. Top enriched clusters are identified on the network. Bar graphs represent the raw P-values for the most enriched GO-terms by (**B**) molecular function and (**C**) biological process.

### SHFL Influences P-body and Stress Granule Dynamics during KSHV infection

Cytoplasmic RNA-granules, such as Stress Granules (SG) and Processing Bodies (P-Bodies), are membrane-free, phase-separated ribonucleoprotein (RNP) complexes that function in the storage, translational arrest, and/or degradation of RNA in the cytoplasm and nucleus (**41–48**). Given the enrichment of SG components in our mass spectrometry data, we next set out to determine whether SHFL localizes to RNP granules. First, HEK293T cells were transfected with either a mock vector or a SHFL expressing vector. Cells were then fixed, permeabilized, and immunostained for known RNP granule markers including DEAD-Box Helicase 6 (DDX6) and enhancer of mRNA decapping 4 (EDC4) for P-bodies and G3BP Stress Granule Assembly Factor 1 (G3BP1) and Cytotoxic Granule Associated RNA Binding Protein (TIA-1) for Stress Granules. RNP granule quantification was performed using CellProfiler to analyze immunofluorescence images stained for the hallmark P-body and SG resident proteins as described in the methods section. In HEK293T cells, SHFL remains diffusely cytoplasmic as we observed previously (**16**) (**Figure 3A**). However, surprisingly, we observed that SHFL expression drastically restricted the number of DDX6 (**Figure 3A**) and EDC4 (**Figure 3B**) puncta per cell relative to mock transfection. Furthermore, we also observed that there was a simultaneous induction of “SG-like densities” co-localizing with SHFL in these same expressing cells (**Figure 3B**). A similar effect was found with TIA-1, a second SG marker, in SHFL expressing cells (**Supp Figure 4**).

**Figure 3:**
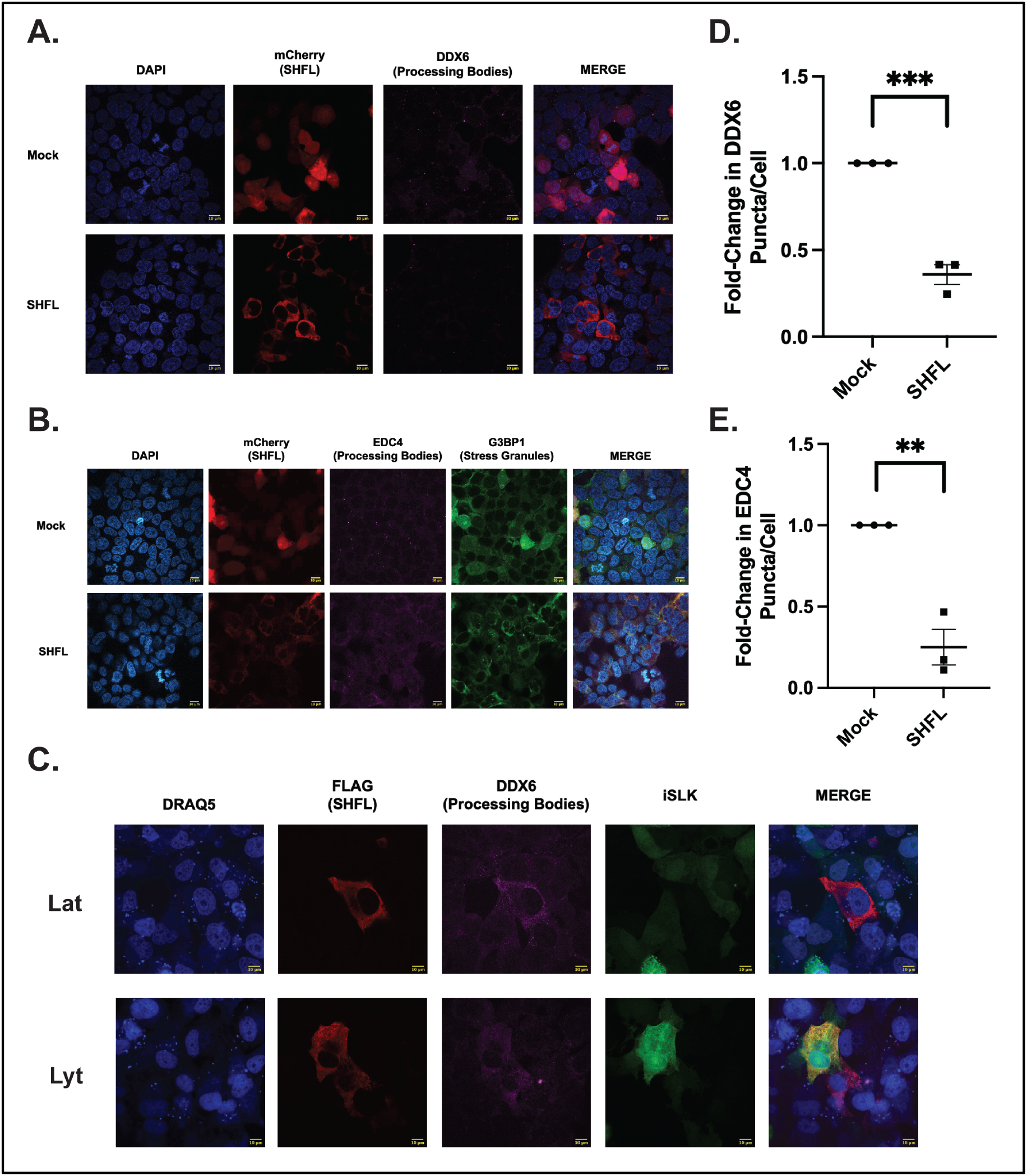
SHFL expression influences RNA granule formation. (**A-B**) HEK293T cells were transfected with either NC-SHFL or a mCherry (mock) only vector. Cells were subjected to immunofluorescence assay and stained for the indicated processing body (DDX6, EDC4, purple) or stress granule markers (G3BP, green). (**C**) iSLK.WT cells were transfected with either a FLAG-tagged SHFL or FLAG-empty (mock) vector and left latent (lat) or reactivated (lyt) for 48h. Cells were then subjected to immunofluorescence assay and stained with the indicated antibodies. (**D-E**) The number of P-body per cell was quantified using the CellProfiler pipeline as described in the methods and normalized to the mock control within each replicate. Scale bar represents 10 μm. Statistics were determined using students paired t-test between control and experimental groups; error bars represent standard error of the mean; n=3 independent biological replicates. n.s.=not significant, ***, P<001.

To determine whether SHFL expression also influences P-body formation during KSHV infection, we next transfected iSLK.WT cells with SHFL, left them either latent or reactivated with Doxycycline and Sodium Butyrate, and stained for the same RNP granule markers. As reported in previous literature, we observed no impact on P-body formation in KSHV latent cells and a corresponding decrease in P-body numbers upon lytic reactivation from latency (**Supp Figure 5**) (**49**). In-line with our observations in HEK293T cells, SHFL expression in both KSHV latent and reactivated cells appears to trigger the P-bodies disassembly (**Figure 3C**). Interestingly, we did not observe a corresponding decrease in DDX6 protein expression in SHFL expressing iSLK.WT cells (**Supp Figure 6**), suggesting that SHFL expression may impact the expression of other P-body scaffolding factors or triggers their relocalization. In summary, our data suggest that SHFL may have more global influence over cellular gene expression including perhaps both host and viral genes. The induction of stress granule-like densities by SHFL also points toward an impact on global gene translation.

### SHFL Interacts with and restricts the expression of KSHV ORF57

Given SHFL effect on ORF50 expression and its influence over RNP granule dynamics, we were particularly intrigued by its interaction with KSHV ORF57 detected in our mass spectrometry screen. We first confirmed the interaction between SHFL and ORF57 by immunoprecipitation and reverse immunoprecipitation (**Figure 4A**). While SHFL overexpression in iSLK.WT cells resulted in lower expression of ORF57 mRNA (**Figure 1A**), transient co-expression of SHFL and ORF57 in HEK293T cells seems to have no effect on ORF57 mRNA (**Figure 4D**), reinforcing the idea that the effect observed on viral mRNA levels in KSHV positive cells is post-transcriptional and stems from a ORF50-dependent mechanism. We thus hypothesized that SHFL could be influencing the expression of ORF57, which in turn could be a mechanistic underpin to SHFL-mediated translational repression of ORF50. First, we showed that the interaction between ORF57 and SHFL is drastically reduced when the samples are treated with RNAse, suggesting that SHFL and ORF57 may be brought together in a complex around viral and perhaps even host RNAs (**Figure 4A**). Next, we found that SHFL co-expression with ORF57 markedly reduced the amount of ORF57 protein, but not mRNA levels, expressed in HEK293T cells (**Figure 4B-D**) pointing once again to a mechanism beyond viral gene transcription and RNA stability. Lastly, we wondered whether SHFL and ORF57 co-localize, and if so, does this co-localization coincide with SHFL induced SG as shown in **Figure 3**. To test this, we again co-transfected SHFL alongside ORF57 in HEK293T cells and performed an immunofluorescence assay. Cells were then fixed, permeabilized, and immunostained for both ORF57 and the SG marker TIA-1. Interestingly, while ORF57 was predominately nuclear, distinct densities of SHFL appeared to co-localize with densities of ORF57 and TIA-1 in the cytoplasm (**Figure 4E-F**). We also did not see the same level of SG marker accumulation with TIA-1 as previously observed with expression of SHFL alone, indicating that ORF57 may still be dispersing SHFL induced RNA granules. While TIA-1 is a SG marker, the protein itself serves a primary function in orchestrating translational arrest (**50,51**). As such, these data indicate that SHFL may be restricting the translation of ORF57 and therein its expression.

**Figure 4:**
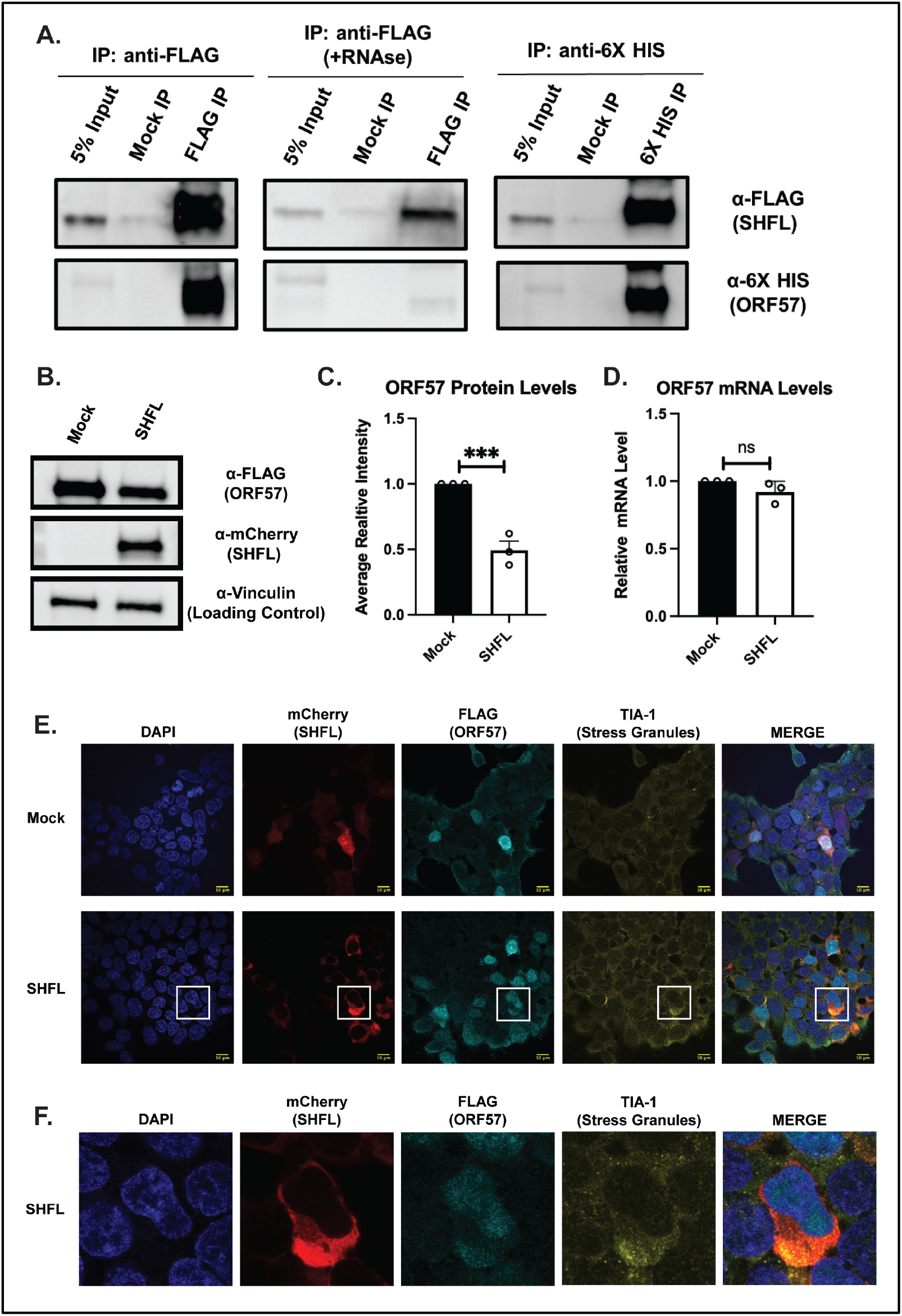
SHFL interacts with and restricts the expression of KSHV ORF57. (**A**) HEK293T cells were co-transfected with both a FLAG-tagged SHFL and a 6X HIS-tagged ORF57. Cells were then harvested, lysed, and co-IP performed using FLAG-tag affinity beads. Reverse co-IP was performed using anti-6X HIS antibody. RNAse co-IP was performed similarly to FLAG co-IP with an additional RNAse A and RNAse T1 treatment prior to overnight co-IP (**B**) HEK293T cells were co-transfected with a FLAG-tagged ORF57 and either a mCherry only (mock) or NC-SHFL vector. Cells were then harvested and subjected to immunoblot and stained with the indicated antibodies (**B**) or total RNA extracted for RT-qPCR to ORF57 mRNA levels (**D**). (**E**) HEK293T cells were transfected with either a mCherry only (mock) or NC-SHFL vector. Cells were subjected to immunofluorescence assay and stained for the indicated proteins. Statistics were determined using students paired t-test between control and experimental groups; error bars represent standard error of the mean; n=3 independent biological replicates. n.s.=not significant, ***, P<001.

## Discussion

Since its initial characterization by Suzuki *et. al*. in 2016, C19ORF66 (here referred to as SHFL), has emerged as a critical piece of the innate immune response to viral infection. SHFL restricts the replication of a vast assortment of DNA, RNA, and retroviruses to varying degrees and is itself an Interferon Stimulated Gene (ISG) (**20,27**). Fascinatingly, for each virus that SHFL restricts, a different mechanism has been described. These inhibition strategies range dramatically from the targeting of viral RNA and protein stability to restriction of viral protein translation, and even the architecture of viral replication organelles (**23**). SHFL is also the first human gene found to actively restrict the −1 programmed ribosomal frameshift (−1PRF), a translation strategy conserved across several eukaryotic viruses, mammals, and even prokaryotes (**19,22,52–54**). Collectively, these studies reflect the versatility of SHFL to broadly influence gene expression and therein directly impact the balance between host and viral gene expression during infection. Similarly, we have previously identified SHFL as a transcript that actively escapes virus-induced RNA decay during KSHV infection (**18**). Furthermore, we also found that the SHFL protein is a potent anti-KSHV factor, restricting nearly every stage of KSHV lytic replication following reactivation from latency. Here, we present our recent efforts toward understanding the mechanism(s) by which SHFL restricts KSHV infection.

Looking broadly at KSHV lytic gene expression, we found that transient SHFL expression stringently restricts all classes of lytic genes at both the RNA and protein levels. Taken together with our previous observations of SHFL knock-down (**18**), this restriction is likely a domino effect from a direct impact of SHFL on KSHV early gene expression. In line with this, we observed a significant restriction of KSHV ORF50 (RTA), which encodes the master latent-to-lytic switch protein responsible for the initiation of the entire lytic gene cascade (**29**). Upon further investigation, we found that SHFL binds to ORF50 mRNA but does not significantly impact the half-life of ORF50 mRNA during lytic replication. This suggests that SHFL influence over ORF50 is likely at the protein level. It would thus be interesting to investigate the effect of SHFL on lysosome and proteasome-mediated degradation during KSHV infection as SHFL has been implicated in these pathways before (**22,25**). This restriction of ORF50 early-on following lytic reactivation would undoubtedly inhibit many crucial stages of KSHV lytic gene expression and therein the remainder of lytic viral replication. Whether SHFL exclusively impacts ORF50 expression during KSHV infection or, more broadly, the protein stability and/or translation of multiple KSHV early genes remains an important future direction for us.

To better understand the mechanism by which SHFL restricts herpesviral translation, we next mapped the interactome of SHFL during KSHV infection using an IP-MS approach. We found an enrichment of cellular RNA-binding proteins (RBPs) that interact with SHFL during both KSHV latency and lytic reactivation. Among these include a vital host translation factor and previously identified SHFL interactor, Polyadenylate-Binding Protein Cytoplasmic 1 (PABPC). We also identified several other RBPs that are known constituents of cytoplasmic RNP granules including both P-bodies and Stress Granules. Both granule types have gained increasing attention over the past decade as critical biophysical sites of RNA regulation that have recently been attribute both pro- and anti-viral functions (**56–59**). Here, we show for the first time that transient expression of SHFL alone triggers the disassembly of P-bodies and simultaneously the induction SG-like densities in virus-free cells. Interestingly, in KSHV-positive cells, we observed a similar reduction in P-bodies during both viral latency and lytic replication but have yet to determine whether an impact can be observed on SG formation. The SG-like densities we have overserved by SHFL suggests that the functions of SHFL likely impact global cellular translation more broadly than initially anticipated, perhaps restricting the translation of several host genes which could in turn influence P-body assembly and/or scaffolding factor localization. These dynamics between SHFL, a known ISG, and RNA granules raises several questions regarding the anti-viral capacity of P-bodies during KSHV infection such as 1) Do P-bodies also have an unforeseen pro-viral role given their disassembly by SHFL, 2) Are transcripts targeted by SHFL localized to SHFL-induced SGs for translational arrest and 3) What part of the SHFL mechanism is responsible for its influence over RNP granules? Moving forward we are interested in exploring the breadth of both viral and cellular transcripts that targeted by SHFL in KSHV infected cells.

The relationship between KSHV and RNA granules is multilayered. While P-bodies are constitutively formed in cells, SG only form in-response to cellular stressors such as oxidative stress and, relevantly, viral infection. P-body dynamics during KSHV latency remains an active area of research to better understand what viral and host factors regulate their stability (**49,60,61**). However, during lytic replication, it has been clearly established that KSHV actively disassembles P-bodies and restricts the formation of SGs. This restriction of RNP granules during lytic replication is directly facilitated by the viral protein KSHV ORF57 (**49,62**). ORF57 is a master regulator of KSHV viral RNA fate with roles ranging from viral mRNA splicing, nuclear mRNA export, and even facilitation of viral mRNA translation in the cytoplasm (**39,63,64**). As such, we were keenly interested in understanding the relationship between SHFL and ORF57. Here, we found that SHFL does in-fact interact with ORF57 in an RNA-dependent manner. Furthermore, we also observed a down regulation of ORF57 expression when co-expressed alongside SHFL. Notably, there was no impact of SHFL on ORF57 mRNA levels, which in combination with our observations of ORF50, further suggests that SHFL also targets ORF57 expression at the protein level. Given its ability to influence RNP granules, we next investigated whether ORF57 could restrict the formation of SHFL induced SG-like densities. Surprisingly, we found that SG were still restricted by ORF57 despite its distinct downregulation by SHFL. In their place, we observed distinct accumulations of ORF57, SHFL, and the SG marker TIA-1 in the cytoplasm. These SHFL-ORF57 densities could reflect sites of translational arrest by SHFL on ORF57 specifically. However, it may also suggest that there could be a distinct difference in the RNP composition of SHFL induced SGs that could be tailored toward a response to viral genes versus host genes. Further exploration is required to determine if SHFL also restricts the translation of ORF57. Or, as suggested by recent SHFL studies in ZIKV, PEDV, and JEV, SHFL could be coordinating with lysosomal or ubiquitinoylation pathways to degrade ORF57 (**22,25,26**).

In conclusion, our findings lay the foundation for a complex relationship between SHFL, a potent anti-viral factor and the DNA virus, KSHV. Following the escape of SHFL mRNA from SOX cleavage, SHFL protein levels climb over the course of KSHV lytic replication. SHFL expression restricts both KSHV early and delayed early genes in a manner that cascades out to late gene expression, an effect whose ramifications are evident across every step of KSHV lytic replication. Here, we show for the first time that overexpression of SHFL influences the formation of cytoplasmic RNA granules, namely stress granules and processing bodies. Thus, SHFL may be restricting herpesviral gene expression at a stage between viral protein stability and viral gene translation. This impact on RNP granules also suggests that SHFL mechanism of action has much broader repercussions on global cellular translation. And therein, SHFL could be restricting the translation of host gene that serve pro-viral roles in the earliest stages of lytic replication. Among the interactions of SHFL, ORF57 also represents a cornerstone of lytic gene expression initiation and countless roles in the stability of viral mRNAs during infection. As such, SHFL could also target ORF57 in a similar manner as ORF50, and through this two-pronged assault, cripple KSHV replication. By studying the impact of SHFL on KSHV lytic reactivation, we continue to unravel unexpected relationships between the regulation cellular RNA fate and the virus-host arms race for control of global gene expression during herpesviral infection.

## Materials and Methods

### Cells and transfections

HEK293T cells (ATCC) were grown in Dulbecco’s modified Eagle’s medium (DMEM; Invitrogen) supplemented with 10% fetal bovine serum (FBS). The KSHV-infected renal carcinoma human cell line iSLK.BAC16 (iSLK.WT) (kind gift from Dr. B. Glaunsinger) bearing doxycycline-inducible RTA was grown in DMEM supplemented with 10% FBS (**65,66**). KSHV Lytic reactivation was induced by the addition of 1 μg/ml doxycycline (BD Biosciences) and 1 mM sodium butyrate for 48 hr as reported above. For DNA transfections, cells were plated and transfected after 24 h when 70% confluent using PolyJet (SignaGen). For small interfering RNA (siRNA) transfections, cells were reverse transfected in 6-well plates by INTERFERin (Polyplus Transfection) with 10 μM siRNAs. siRNAs were obtained from IDT as Dicer-substrate siRNA (DsiRNA; siRNA C19ORF66, hs.Ri.C19orf66.13.1).

### Plasmids

The C19ORF66 coding region was obtained as a gBlock from IDT and cloned in a pcDNA4 Nter-3×Flag vector (FLAG-SHFL). The SHFL coding region was then cloned into a pmCherry-C1 vector (kind gift from Jeffrey Kane) to construct the N-terminal mCherry-C19ORF66 vector (NC-SHFL). KSHV lytic gene coding sequences (ORF50, ORF57, ORF59, and ORF52) were derived from a library of strep-tagged KSHV ORFs (Davis et. al) and cloned into recipient vectors to construct ORF57-6XHIS and FLAG-ORF57.

### RT-qPCR

Total RNA was harvested using TRIzol according to the manufacture’s protocol. cDNAs were synthesized from 1 μg of total RNA using AMV reverse transcriptase (Promega) and used directly for quantitative PCR (qPCR) analysis with the SYBR green qPCR kit (Bio-Rad). Signals obtained by qPCR were normalized to those for 18S unless otherwise noted. Primers used in the study are listed in **Supp Table 3**.

### Immunoblotting

Cell lysates were prepared in lysis buffer (NaCl, 150 mM; Tris, 50 mM; NP-40, 0.5%; dithiothreitol [DTT], 1 mM; and protease inhibitor tablets) and quantified by Bradford assay. Equivalent amounts of each sample were resolved by SDS-PAGE and immunoblotted with each respective antibody at 1:1,000 in TBST (Tris-buffered saline, 0.1% Tween 20). Antibodies used for human and KSHV targets are listed in **Supp Table 4**. Primary antibody incubations were followed by horseradish peroxidase (HRP)-conjugated goat anti-mouse or goat anti-rabbit secondary antibodies (1:5,000; Southern Biotechnology).

### Immunoprecipitation

Cells were lysed in low-salt lysis buffer (150 mM NaCl, 0.5% NP-40, 50 mM Tris [pH 8], 1 mM DTT, and protease inhibitor cocktail), and protein concentrations were determined by Bradford assay. At least 400 μg of total protein were incubated overnight with the designated antibody and then with protein G-coupled magnetic beads (Life Technologies) for 1 h. For FLAG construct pull-downs, total protein lysates were instead incubated overnight with Anti-FLAG M2 Magnetic Beads (Sigma) or G-coupled magnetic beads. Beads were then washed extensively with lysis buffer. Where indicated, samples were treated with both RNAse A and T1 for 15 min at RT prior to immunoprecipitation. Lastly, samples were resuspended in 4X laemmli loading dye before resolution by SDS-PAGE.

### Mass Spectrometry

Briefly, iSLK.WT cells were seeded into 10-cm plates and reactivated at the same time. Following 72 Hours post-reactivation, cells were harvested and lysed, and immunoprecipitation for SHFL was performed overnight at 4C. Samples were extensively washed, and trypsin digested overnight. Samples were then cleaned up using a C_18_ column and mass spectral data obtained from the University of Massachusetts Mass Spectrometry Center using an Orbitrap Fusion mass spectrometer. Raw data was filtered based on the number of peptides for each hit and Gene Ontology (GO) enrichment analysis was performed on the human interacting proteins of SHFL using DAVID bioinformatic database. Top enriched clusters are identified on the network.

### RIP

Following transfections, cells were crosslinked in 1% formaldehyde for 10 minutes, quenched in 125mM glycine and washed in PBS. Cells were then lysed in low-salt lysis buffer [NaCl 150mM, NP-40 0.5%, Tris pH8 50mM, DTT 1mM, MgCl2 3mM containing protease inhibitor cocktail and RNase inhibitor] and sonicated. After removal of cell debris, Anti-FLAG M2 Magnetic Beads or Magnetic G-coupled beads were added as indicated overnight at 4°C. The following day, beads were washed three times with lysis buffer and twice with high-salt lysis buffer (low-salt lysis buffer except containing 400mM NaCl). Samples were then separated into two fractions. Beads containing the fraction used for immunoblotting were resuspended in 30μL lysis buffer. Beads containing the fraction used for RNA extraction were resuspended in Proteinase K buffer (NaCl 100mM, Tris pH 7.4 10mM, EDTA 1mM, SDS 0.5%) containing 1μL of PK (Proteinase K). Samples were incubated overnight at 65°C to reverse crosslinking. Samples to be analyzed by immunoblot were then supplemented with 10μL of 4X loading buffer before resolution by SDS-PAGE. RNA samples were resuspended in Trizol, and total RNA was extracted for qPCR as described above.

### RNA Half-life

iSLK.WT cells were plated and transfected with either a mock or NC-SHFL vector. Twenty-four hours later, KSHV was reactivated as described above for roughly 40 hrs. Eight hours prior to the 48 hr timepoint, transfected cells were treated with 5 ug/mL of Actinomycin D to inhibit cellular transcription and cells were collected at the indicated time points from 0-8 hrs. Total RNA was extracted from all samples and to qPCR analysis using primers targeting KSHV ORF50 and normalized to the level of GAPDH mRNA.

### Immunofluorescence

293T or iSLK.WT cells were grown on coverslips and fixed in 4% formaldehyde for 20 min at room temperature. Cells were then permeabilized in 1% Triton X-100 and 0.1% sodium citrate in phosphate-buffered saline (PBS) for 10 min, saturated in bovine serum albumin (BSA) for 30 min, and incubated with the designated antibodies at various dilutions (refer to **Supp Table 2**). After 1 h, coverslips were washed in PBS and incubated with Alexa Fluor 680, 594 or 488 secondary antibodies at 1:1,500 (Invitrogen). Coverslips were washed again in PBS and mounted in DAPI-containing Vectashield mounting medium (Vector Labs) to stain cell nuclei before visualization by confocal microscopy on a Nikon A1 resonant scanning confocal microscope (A1R-SIMe). The microscopy data were gathered in the Light Microscopy Facility and Nikon Center of Excellence at the Institute for Applied Life Sciences, UMass Amherst, with support from the Massachusetts Life Sciences Center.

### RNA Granule Quantification

Processing bodies and stress granules were quantified using an unbiased image analysis pipeline generated in the freeware CellProfiler (cellprofiler.org) (**61,66,67**). First, detection of nuclei in the DAPI channel image was performed by applying a binary threshold and executing primary object detection between 50 and 250 pixels. From each identified nuclear object, the “Propagation” function was performed on the respective 594 channel (denoting Mock (mCherry) or NC-SHFL) image to define transfected cell borders. The identified cell borders were masked with the identified nuclei to define a cytoplasm mask. The cytoplasm mask was then applied to the processing body/stress granule puncta channel images (stains with DDX6 and EDC4 for processing bodies) to ensure only cytoplasmic puncta were quantified. Background staining was reduced in the cytoplasmic puncta channel using the “Enhance Speckles” function. Using “global thresholding with robust background adjustments”, puncta within a defined size and intensity range were quantified. Size and intensity thresholds were unchanged between experiments with identical staining parameters. Intensity measurements of puncta were quantified. Quantification data was exported and used for data analysis.

### Statistical analysis

All results are expressed as means ± standard errors of the means (SEMs) of experiments independently repeated at least three times. Unpaired Student’s *t* test was used to evaluate the statistical difference between samples. Significance was evaluated with *P* values as follows: * p<0.05; ** p<0.01; *** p<0.001.

## Acknowledgments

We thank all members of the Muller lab for helpful discussions and suggestions. Special thanks to Dr. Britt Glaunsinger for both our iSLK cell lines and KSHV antibodies. We are also particularly grateful to Dr. Jennifer Corcoran and Beth Castle for our conversations and suggestions with RNA granules detection and quantification. We would also like to thank Dr. James Chambers at the UMASS IALS Light Microscope Facility and Dr. Stephen Eyles at the UMASS IALS Mass Spectrometry facility for their help with protocol development and data acquisition.

## Figure Legends

**Supplementary Figure 1: SHFL Broadly restricts KSHV Lytic Gene Expression**. (A) Full representation of RT-qPCR data summarized by heat map in Figure 1A. iSLK.WT cells were transfected with a FLAG-tagged SHFL or a FLAG-empty vector and reactivated with doxycycline and sodium butyrate for 48h. Total RNA was then harvested and subjected to RT-qPCR to measure mRNA levels of the indicated viral early, delayed early, and late genes. Statistics were determined using students paired t-test between control and experimental groups; error bars represent standard error of the mean; n=3 independent biological replicates. ***, P<001.

**Supplementary Figure 2: ORF50 mRNA Stability Assays**. iSLK.WT cells were transfected with a FLAG-tagged SHFL or a FLAG-empty vector and reactivated with doxycycline and sodium butyrate for 48h. 6h prior to the 48h mark, cells were treated with 5 ug/ml Actinomycin D and harvested over an 8h time course. (A) Total RNA was then harvested and subjected to RT-qPCR to measure ORF50 mRNA levels. (B) Cells were also harvested, lysed, and total protein extracted and subjected to immunoblot and stained with the indicated antibodies. Statistics were determined using students paired t-test between control and experimental groups; error bars represent standard error of the mean; n=3 independent biological replicates. N.S., not significant.

**Supplementary Figure 3: SHFL pull-down for IP-MS and PABPC Co-IP**. iSLK.WT cells were reactivated using doxycycline and sodium butyrate for 72h. Cells were then harvested, lysed, and a co-IP was performed using the C19ORF66 (SHFL) antibody. Samples were then subjected to immunoblot and stained with the indicated antibodies.

**Supplementary Figure 4: SHFL expression induces Stress granule-like densities in HEK293T cells**. HEK293T cells were transfected with either NC-SHFL or a mCherry vector. (A) HEK293T cells were transfected with either NC-SHFL or a mCherry (mock) only vector. Cells were then subjected to immunofluorescence assay and stained for the indicated stress granule markers (G3BP, green and TIA-1, yellow). (B-C) Quantification of SG markers in transfected cells using CellProfiler as described in the methods.

**Supplementary Figure 5: DDX6 expression in SHFL transfected HEK239T and iSLK.WT cells. (**A) iSLK.WT cells were transfected with either NC-SHFL or a mCherry only vector and reactivated with Doxycycline and Sodium Butyrate for 48h. Cells were then harvested, lysed, and total protein extracts and subjected to immunoblot and stained for the indicated antibodies.

**Supplementary Figure 6: P-body formation during KSHV latency and lytic replication**. iSLK.WT cells left latent (lat) or reactivated (lyt) with Doxycycline and Sodium Butyrate for 48h. Cells were then subjected to immunofluorescence assay and stained for the indicated P-body marker (DDX6, magenta)

**Supplementary Table 1: List of SHFL Interactors during KSHV infection**.

**Supplementary Table 2: DAVID Go-term analysis of human SHFL interactors**.

**Supplementary Table 3: List of Antibodies used in this study**.

**Supplementary Table 4: List of Primers used in this study**

