## Supplementary figures and images for "Shiftless Restricts Viral Gene Expression and Influences RNA Granule Formation during KSHV lytic replication"

### Supp Figure 1

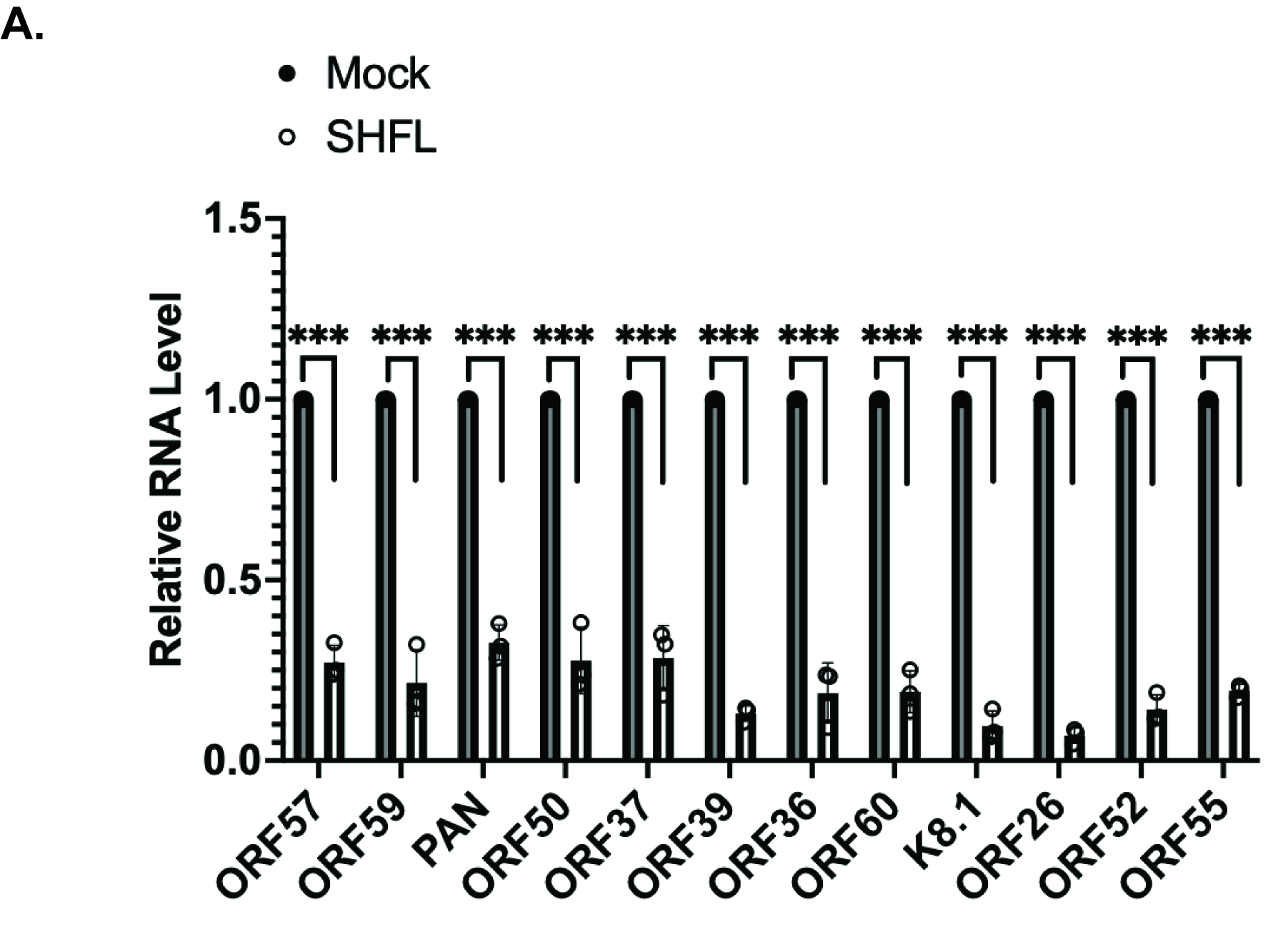

### Supp Figure 2

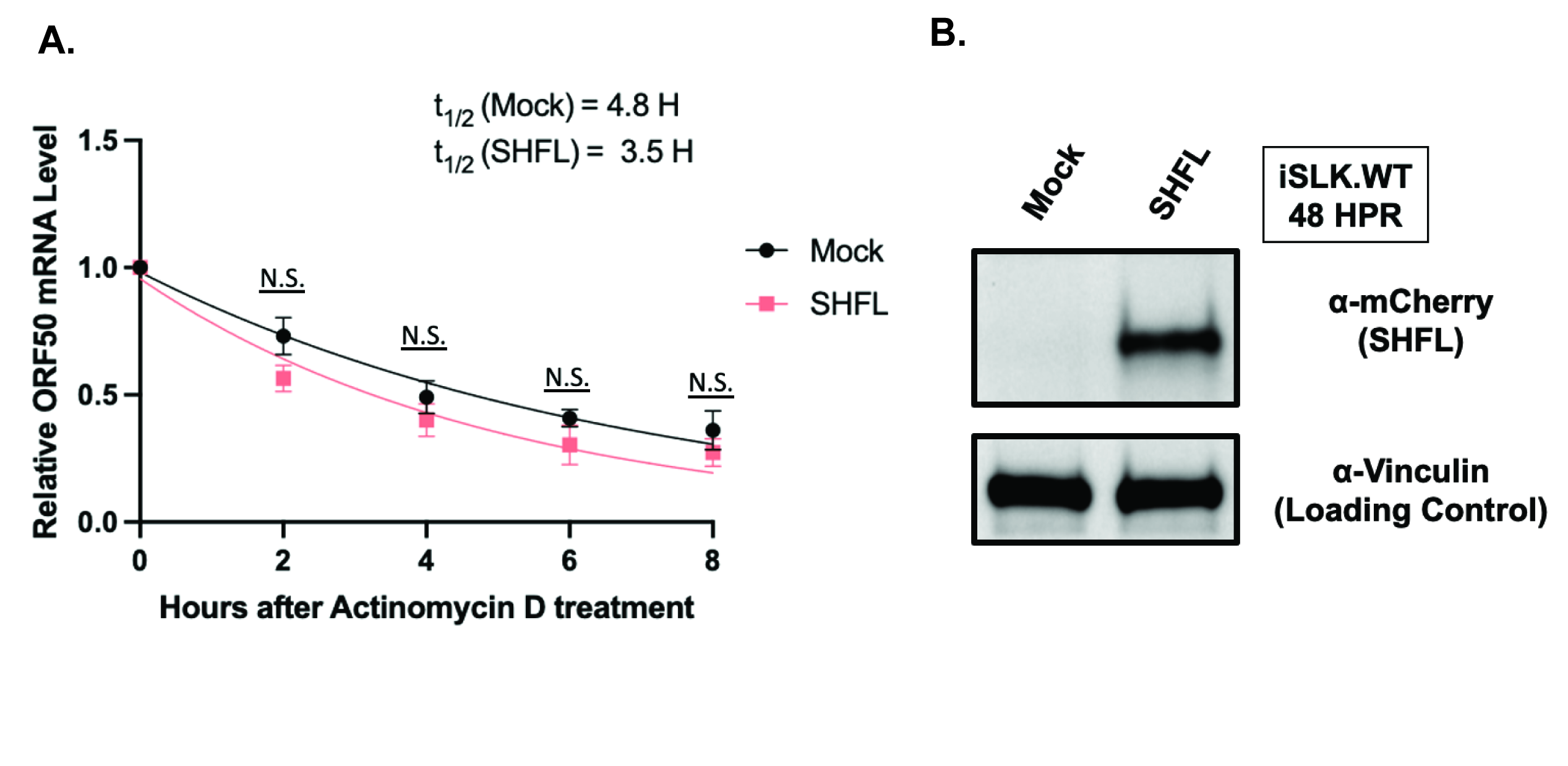

### Supp Figure 3

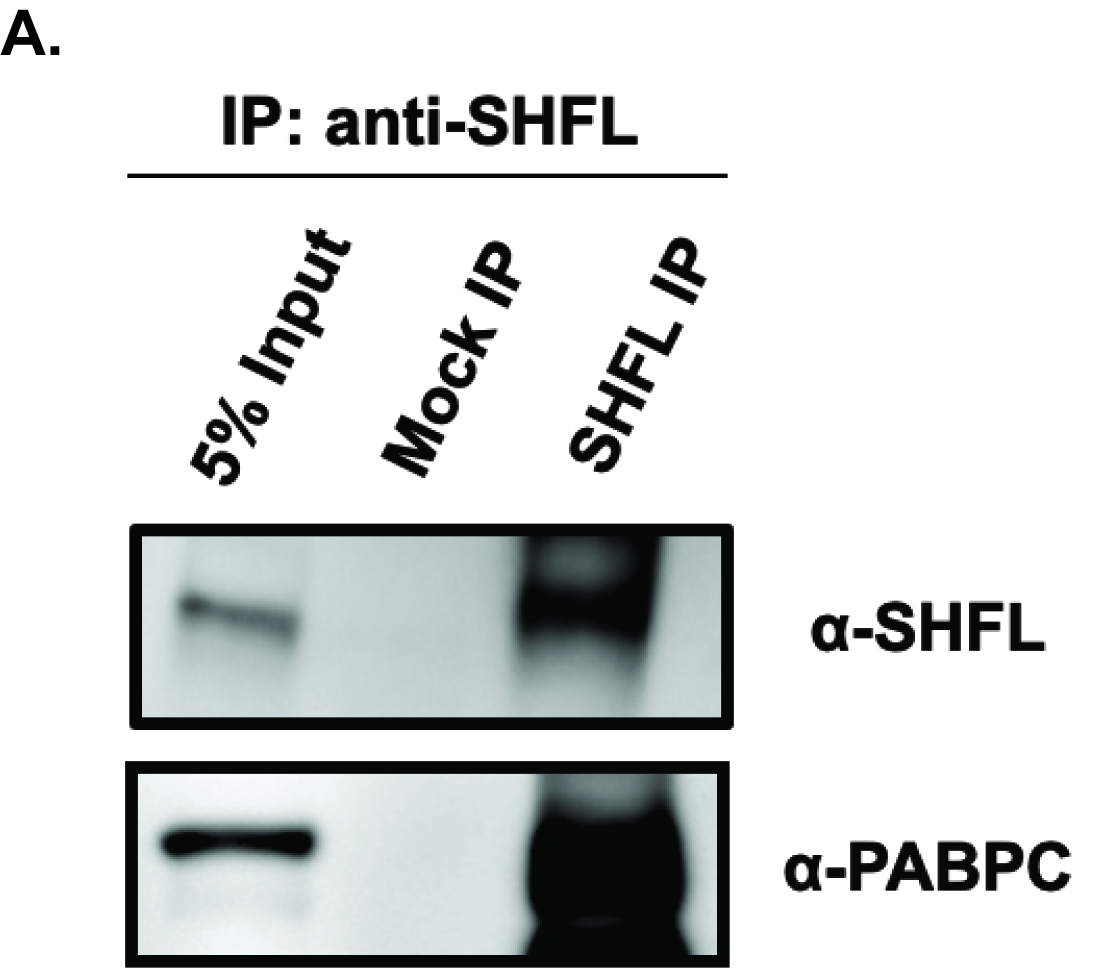

### Supp Figure 4

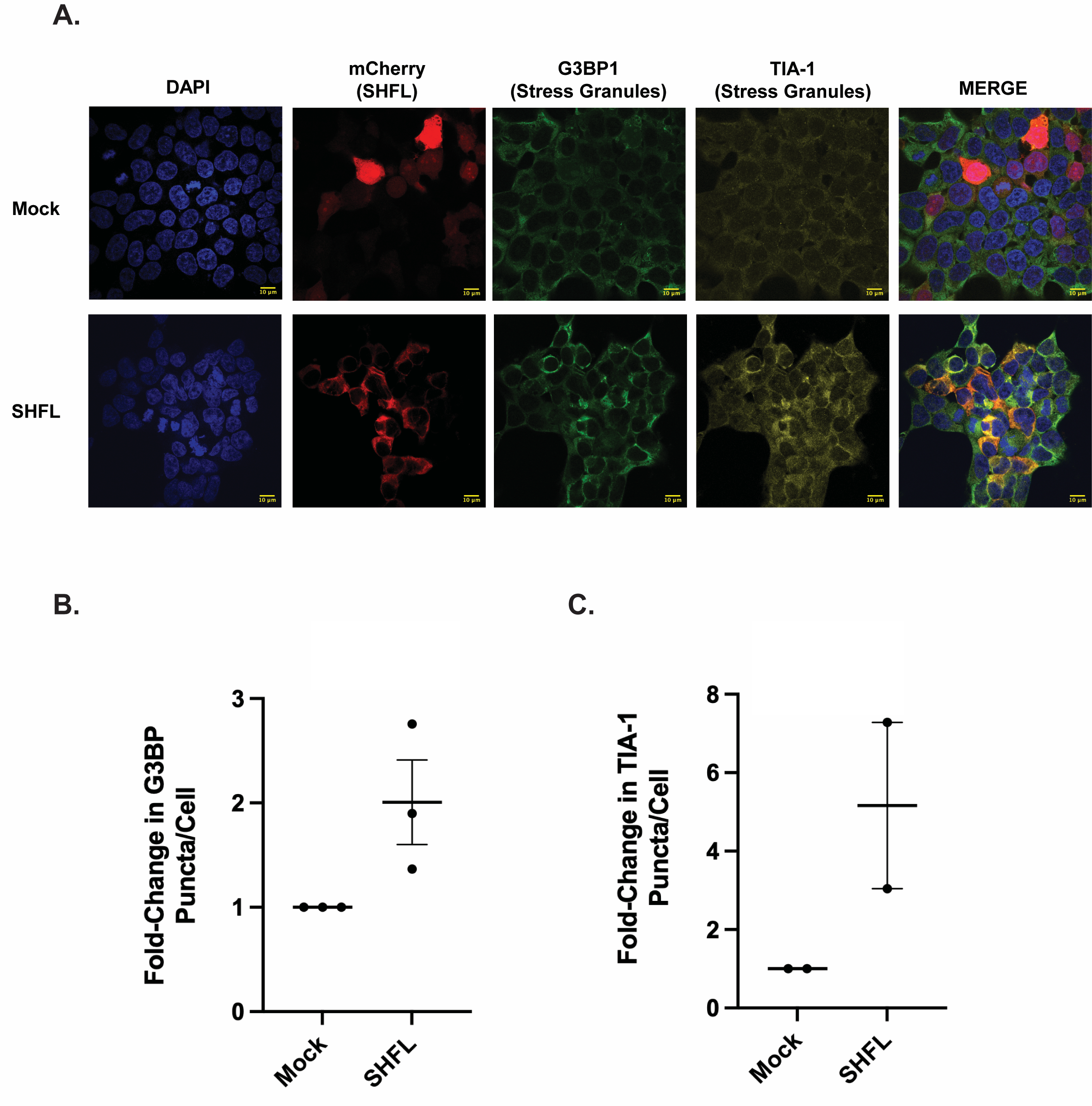

### Supp Figure 5

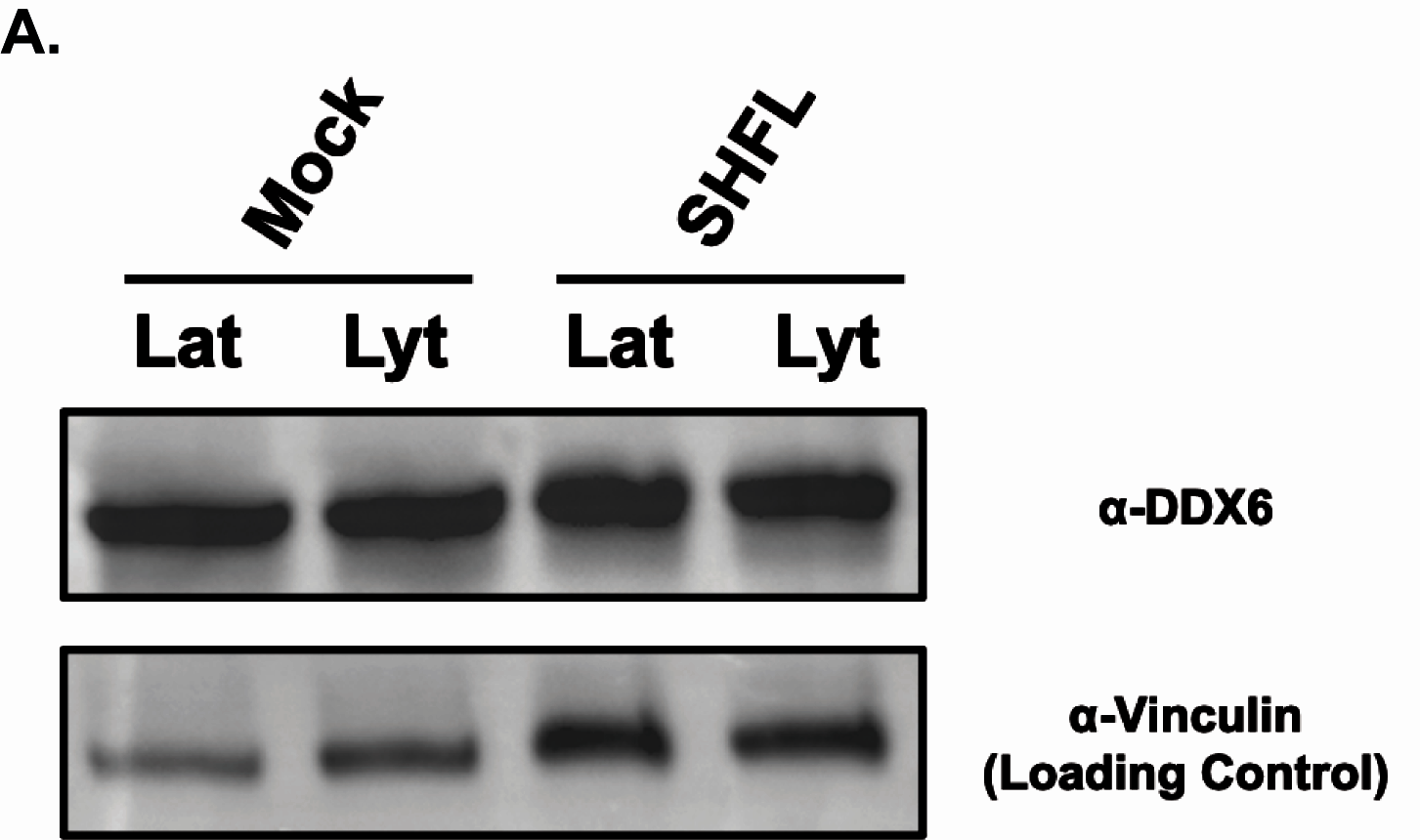

### Supp Figure 6

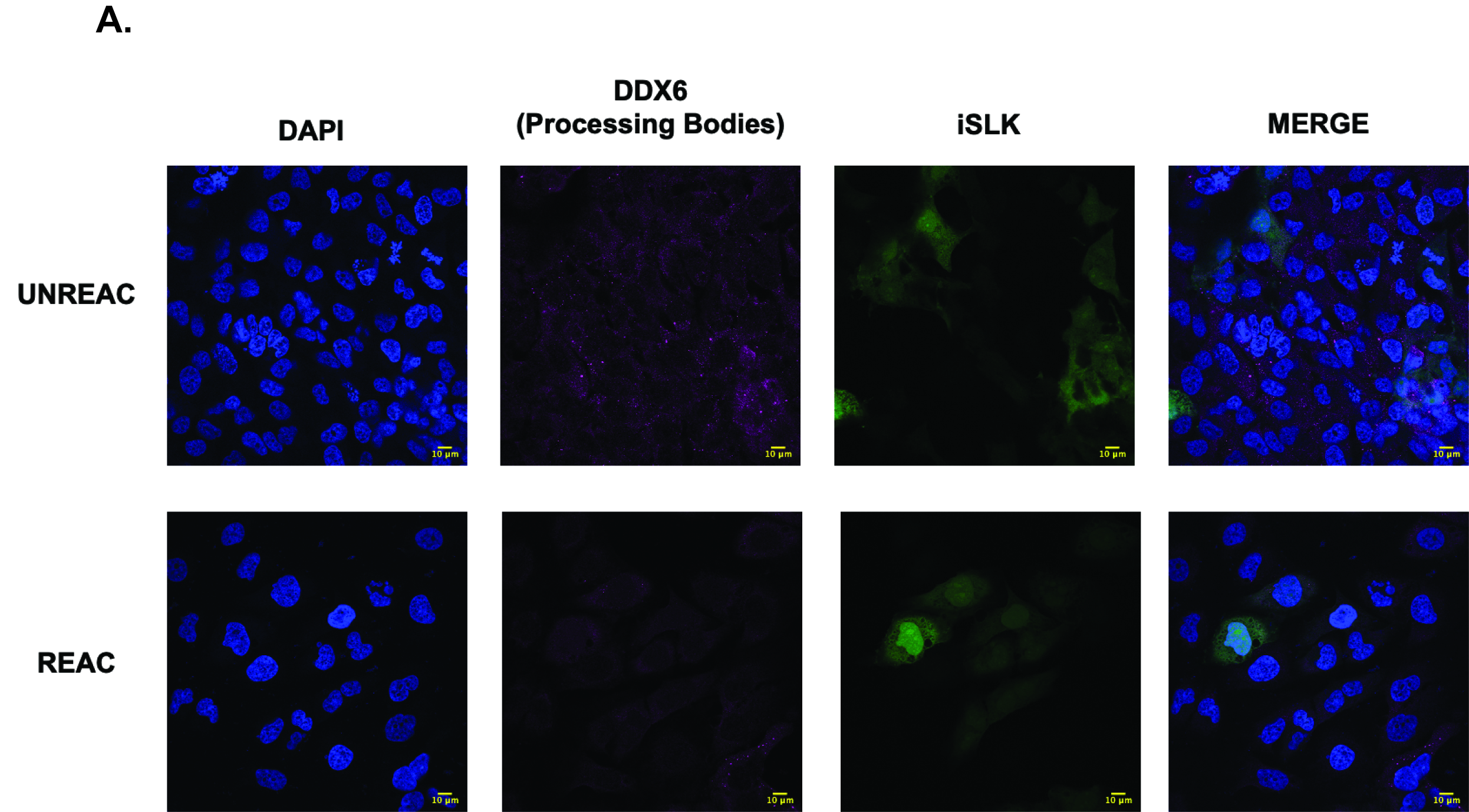
